# sRNA-mediated crosstalk between cell wall stress and galactose metabolism in *Staphylococcus aureus*

**DOI:** 10.1101/2024.11.18.624069

**Authors:** Maëliss Germain, Hugo Robin, Kim Boi Le Huyen, Sébastien Massier, Nicolas Nalpas, Julie Hardouin, Philippe Bouloc, Astrid Rouillon, Svetlana Chabelskaya

**Affiliations:** Inserm, BRM [Bacterial RNAs and Medicine] - UMR_S1230, 35033 Rennes, France; University of Rouen Normandie, INSERM US 51, CNRS UAR 2026, HeRacLeS-PISSARO, Normandie University, 76000 Rouen, France; University of Rouen Normandie, INSA Rouen Normandie, CNRS, Polymers, Biopolymers, Surfaces Laboratory UMR 6270, F-76000 Rouen, France; University Paris-Saclay, CEA, CNRS, Institute for Integrative Biology of the Cell (I2BC), 91198 Gif-sur-Yvette, France

## Abstract

*Staphylococcus aureus* is an opportunistic pathogen responsible for a wide range of diseases in humans. During infections, this bacterium is exposed to various stresses that target its cell wall, such as oxidative or acid environments as well as various cell wall-acting antimicrobials. *S. aureus* has effective regulatory systems for responding to environmental stresses enabling the expression of factors necessary for its survival. Bacterial small RNAs (sRNAs) play a crucial role in this adaptation process. In this study, we show that RsaOI, an *S. aureus* sRNA, accumulates under acid stress conditions. This response is mediated via the two-component system VraSR, which is associated with the cell wall damage response. As a component of the VraSR regulon, RsaOI contributes to the survival of *S. aureus* under acid stress and affects its susceptibility to glycopeptide antibiotics. Our findings reveal that RsaOI targets *lacABCDFEG* operon, which encodes components of tagatose pathway, a unique mechanism responsible for galactose metabolism in *S. aureus*. By antisense base pairing near the ribosome-binding site of *lacD*, RsaOI inhibits the expression of this gene, encoding tagatose-6-phosphate aldolase. This regulation disrupts the tagatose pathway, impairing galactose utilization in *S. aureus*. These findings highlight the role of RsaOI in the mediation between cell wall stress responses and specific metabolic pathway.

## INTRODUCTION

*Staphylococcus aureus* is a member of the commensal human microbiota that can act as an opportunistic pathogen, responsible for both nosocomial and community-acquired infections (1). Its remarkable ability to cause a wide range of infections and establish successful colonization is largely due to its adaptability and resilience. *S. aureus* can survive and proliferate in various hostile environments within the human host, including acidic conditions (2). Such acidic microenvironments are commonly encountered on mucosal surfaces, in infected tissues where the immune response creates localized acidic conditions, and within phagolysosomes in the case of intracellular bacteria. To withstand these stresses, *S. aureus* employs a variety of adaptive mechanisms, including regulatory networks that allow rapid modulation of cellular processes essential for survival (2). One such adaptation involves reducing the permeability of the cell membrane and cell wall to protons by altering their charge composition. During acid stress, Gram-positive bacteria increase the expression of genes involved in cell wall modifications, such as the *cap* gene, linked to capsule biosynthesis, and the *dlt* operon, which adds positive charges to teichoic acids, enhancing resistance to acidic conditions (3–5). Another approach is the use of proton pumps such as the F_0_F_1_-ATPase, which pump excess protons out of the cell (6, 7). Additionally, bacteria increase the concentration of alkaline compounds within the cell to counteract the acidification of the cytoplasm. For this, *S. aureus* increases the import of amino acids and osmolytes to maintain intracellular pH through decarboxylation reactions that consume protons (8, 9). Moreover, ammonia production is also promoted, either from urea through urease activity or via the arginine deaminase system (7, 10). Finally, energy metabolism and metabolic pathways are significantly reoriented. Respiration is favoured over fermentative metabolism to limit additional acidification of the environment (8).

Global transcriptomic analyses have offered insights into how *S. aureus* adjusts its gene expression profiles to adapt to acidic environments, revealing the complexity of its stress response mechanisms (5, 7, 11, 12). Despite these advances, the regulators that coordinate acid survival mechanisms remain partially characterized. Studies focused on a small number of transcriptional regulators provide only fragmentary insights into the network, yet they hint at its complexity. For instance, transcription of urease, a crucial enzyme in acid stress responses, was shown to be controlled by global regulators CcpA, CodY, and Agr (13). In addition, two-component systems (TCS), which integrate a signaling sensor kinase and a response regulator, play important roles in adapting to acidic stress. For example, the GraRS TCS has been shown to be involved in resistance to antimicrobial peptides under acidic conditions (14) and to enhance *S. aureus* survival within the acidified phagolysosomes of macrophages (15). In addition, other staphylococcal TCS, such as KdpDE and VraSR, were also proposed to participate in the control of low pH survival (2, 8).

Although transcriptomic studies provide extensive information on acid stress responses, they often overlook post-transcriptional regulation. However, adapting to environmental changes often requires transcription factors to work in conjunction with post-transcriptional mechanisms mediated by small RNAs (sRNAs) (16, 17). Most sRNAs are RNA molecules between 50 and 500 nucleotides in length and play crucial roles in various cellular processes, especially under specific growth and stress conditions, enabling faster regulatory responses than transcriptional mechanisms alone (18). Typically, sRNAs act through antisense base pairing with the mRNA of their target genes (18–20). While numerous studies highlight the involvement of diverse protein regulators, the role of sRNAs in orchestrating these adaptive responses remains less understood, despite emerging evidence suggesting that sRNAs are essential for fine-tuning responses to environmental stressors.

In this study, we analyzed transcriptomic responses of *S. aureus* under acidic pH, with a focus on sRNA expression. Our analysis showed that a notable sRNA, RsaOI (Srn1490_RsaOI_Teg47)(21), exhibits a marked increase in expression under acidic conditions. We further investigated the regulatory mechanisms governing RsaOI expression and characterized its targetome. Our data reveal that the TCS VraSR, known for its role in cell wall stress response, is activated under acidic conditions and induces the expression of RsaOI. RsaOI targetome analysis identified that this sRNA inhibits the expression of *lacD.* The latter is part of the lactose operon that is responsible for lactose and galactose metabolism. These findings suggest that RsaOI functions as a regulatory mediator, linking cell wall stress responses with metabolic adjustments, particularly in coordinating lactose metabolism under acid stress. This crosstalk highlights a regulatory strategy by which *S. aureus* fine-tunes its metabolic priorities to optimize growth and survival in challenging environments.

## MATERIAL & METHODS

### Bacterial strains, plasmids and growth conditions

The strains and plasmids utilized in this study are detailed in Supplementary Table S1. Supplementary Table S2 lists all the primers used. Unless otherwise specified, *S. aureus* strains were grown in Brain Heart Infusion medium (BHI, Oxoid) at 37°C with shaking (160 rpm) or on BHI agar (Oxoid) at 37°C. *Escherichia coli* DH5-α strains were grown in LB (lysogeny broth) at 37°C with shaking (160 rpm) or on LB agar plates at 37°C. For the maintenance of plasmids or resistance cassettes, antibiotics were used at the following concentrations: for *S. aureus*, erythromycin and chloramphenicol at 10 µg/ml; for *E. coli*, ampicillin at 50 µg/ml. The purified plasmids were used for the transformation in *S. aureus* RN4220 strain by electroporation. The φ80 phages prepared from RN4220 were then used to transduce plasmids in HG003 strains. pIMAYΔ*lacR* vector is a pIMAY (22) derivative containing the PCR-amplified *lacR* upstream and downstream sequences cloned by Gibson assembly (using primers from Supplementary Table S2) as described (23). The *lacR* gene was deleted in *S. aureus* HG003 strain using pIMAY as described previously (23). Other *S. aureus* mutants were constructed by transducing the erythromycin resistance cassette from the Nebraska Tn mutant library USA300 (24) into the HG003 strain using φ80 phage.

To induce acidic stress, cells were grown for two or six hours in a LB medium supplemented with 100 mM HEPES to maintain pH to 7. Cells were then divided, pelleted by centrifugation at 3000 rpm for 5 minutes, and resuspended in fresh LB medium buffered with HEPES to pH5 or pH7 for 30 minutes.

For metabolic tests, *S. aureus* strains WT, Δ*rsaOI* (both carrying pICS3 plasmid), or Δ*rsaOI* complemented strain (carrying pICS3-P*amiA-rsaOI* plasmid), were cultured at 37°C in LB or NZM broth and then diluted to 1:100 ratio in fresh media. When necessary, the media were supplemented with glucose or galactose at a concentration of 11 mM, and HEPES at 100 mM. Growths were measured by Biotek microplates reader.

### Construction of *gfp*-reporter transcriptional and translational fusions

pCN33 and pCN38-based vectors containing *gfp-*reporter fusions were constructed using primers from Supplementary Table 2. *gyrB* sequence of pCN33-*PtufA-gyrB-gfp* vector was replaced by 5’ *lacD* sequence to produce pCN33-*PtufA-lacD-gfp.* For the pCN38-*PrsaOI-gyrB-gfp* construction, *rsaOI* promoter region (−99 nucleotides to +39 after the *rsaOI* transcriptional start) was amplified from *S. aureus* HG003 genomic DNA by PCR. The *rsaOI* promoter fragment was placed instead of P*tufA* promoter of vector pCN33-*PtufA-gyrB-gfp*. All cloning experiments were performed with Gibson Assembly Master Mix (New England Biolabs). The reactions were then transformed into *E. coli* DH5-α by heat shock at 42°C. The constructs were confirmed by DNA sequencing, and then electroporated into RN4220 before being transduced into HG003 or its derivatives using bacteriophage Φ80. When necessary, *S. aureus* HG003 strain was used to co-transform the *lacD-gfp* fusion vector with the *rsaOI* expressing plasmid. Cultures of these co-transformed *S. aureus* strains were grown at 37°C in LB or BHI supplemented with 10 μg/ml chloramphenicol and erythromycin. Fluorescence and OD_600_ measurements were driven by Biotek microplates reader as previously described (25). To induce *rsaOI* expression, cells were grown in LB medium for two hours, then divided, with one sample receiving 10 µl of HCl 37% per 10 ml of culture.

### Induction of acidic stress for acid susceptibility tests and proteomic studies

To induce *rsaOI* expression by acidic pH, cells were grown for two hours in a LB medium supplemented with 100 mM HEPES to maintain pH at 7. Cells were pelleted by centrifugation at 3000 rpm for 5 minutes and resuspended in a fresh LB medium buffered with HEPES to pH5 for an additional 2 hours. After stress induction, cells were collected and pelleted by centrifugation for proteomic studies or plated on BHI agar plates for CFU counting.

### Antibiotic susceptibility tests

Ten-fold serial dilutions of overnight cultures of WT HG003, Δ*rsaOI* or complemented strain (Δ*rsaOI/p-rsaOI*) were plated and incubated for 24 hours at 37°C on TSA or TSA supplemented with 1 µg/ml vancomycin. Spot population analysis profile assay (spot PAP) allows rapid, sensitive and reproducible qualitative and quantitative testing of antibiotic resistance. For this, five microliters of each dilution were dropped on TSA or TSA supplemented with 1 µg/ml vancomycin, and then incubated for 24°C at 37°C. Dilutions were deposed from top to the most-concentrated (without dilution) to the dilution 10^-6^.

### RNA-sequencing method

Total RNA extraction was performed as previously described (26) on three independent replicates. RNA quality was then evaluated on a High Sensitivity RNA Screen Tape chip (Agilent). rRNA depletion, cDNA library and sequencing were performed by Genewize® platform using their Strand-Specific RNA-seq protocol. The resulting FASTQ raw data were imported into the Galaxy web platform, using the public server at https://usegalaxy.eu/ for data analysis. Reads were cleaned using Trimmomatic (version 0.39) and mapped to the *S. aureus* NCTC8325 reference genome, including the file containing sRNA genes annotated for NCTC8325 (https://srd.genouest.org/browse/NCTC8325), using the BWA (version 0.7.18) algorithm. Counting files were generated with HTseq-count (version 2.0.5) and differentially expressed transcripts were identified using DEseq2 (version 1.40.2). Transcripts and genes with a *p-value* < 0.05 and a fold change > 2 were considered significantly differentially expressed across biological replicates.

### Preparation of samples for proteomic studies

Bacteria were prepared as described in the section on acidic stress induction. After pH induction, samples were centrifuged at 4500 rpm for 15 min at 4°C, and washed twice with cold PBS. The pelleted cells were then resuspended in 500 µl of Tris-HCl (pH8) supplemented with a mini cOmplete (Roche) protease inhibitors cocktail, 2 U of DNAse, 2 U of RNAse. Bacteria were lysed using a Fast-Prep system, then centrifuged for 10 min at 12,000 rpm at 4°C. Protein concentration was measured using a Qubit assay, and 500 µg were precipitate with 17.5% TCA overnight at 4°C. The samples were centrifuged at 9000 rpm for 15 min at 4°C, and supernatant was discarded. The pellets were rinsed twice with 2 volumes of cold acetone (100%), and mixed by inversion, followed by the same centrifugation as described before. Samples were dried in a speed vacuum and stored at −80°C.

### Proteins solubilization and quantification

TCA-precipitated proteins were then solubilized in 200 µL of R2D2 denaturing buffer (7 M urea, 2 M thiourea, tri-N-butylphosphine (TBP) 2 mM, dithiothreitol (DTT) 20 mM, 0.5% (w/v) 3-(4-heptyl)phenyl-3-hydroxypropyl)dimethylammoniopropanesulfonate (C7BzO), 2% (w/v) 3-[(3-cholamidopropyl)dimethylammonio]-1-propanesulfonate hydrate (CHAPS)) and sonicated for 30 sec. Protein concentrations were evaluated by Bradford analysis (Bio-Rad). Samples were stored in aliquots (30 µg) at −20 °C until further use.

### Protein Digestion

Twenty-five micrograms of proteins were mixed with sodium dodecyl sulfate (SDS) loading buffer (62 mM Tris-HCl pH 6.8, 20% glycerol (v/v), 0.04% bromophenol blue (w/v), 0.1 M DTT, SDS 4% (w/v)) heated at 95°C for 5 min, and then loaded onto a large SDS-PAGE stacking gel 7%. An electrophoresis was performed (10 mA, 2 hours) to concentrate proteins. After migration, gels were stained with Coomassie Blue G250 and destained (50% ethanol (v/v) and 10% acetic acid (v/v)). The revealed protein band was excised, completely destained and washed three times with water. Samples were then reduced with 10 mM Dithiothreitol (DTT) for 1 hour at room temperature and alkylated with 15 mM Iodoacetamide for 45 minutes in the dark. Gel bands were treated with 50% acetonitrile (ACN)/50% ammonium bicarbonate 10 mM, pH 8 (2 times, 5 min) and dried with 100% ACN (3 times, 10 min). Then, proteins were digested with trypsin (1 μg per band, ratio of 1:25), overnight at 37 °C, in ammonium bicarbonate 10 mM, pH 8. Peptides were extracted with 100% ACN (3 times, 10 min) and then dried using a Speedvac concentrator (SPD111V, Thermo Fisher Scientific) and stored at −20 °C. For each type of bacteria (wild type and mutants), five biological replicates were carried.

### LC-MS/MS analysis

Peptides were solubilized in 0.1% formic acid (FA) (v/v) and quantified using Pierce quantitative colorimetric peptide assay (Thermo scientific). Peptides (0.2 µg) were subjected to quantitative LC-MS/MS analysis on a high-resolution Orbitrap Eclipse Tribrid mass spectrometer coupled to a Proxeon Easy nLC 1200 (Thermo Scientific). Samples were injected onto an enrichment column (Acclaim PepMap C18, 2 cm × 75 μm, 100 Å, Thermo Scientific). The separation was performed with an analytical column needle (Acclaim PepMap C18, 25 cm × 75 μm, 2 μm, 100 Å, Thermo Scientific). The mobile phase consisted of H2O/0.1% formic acid (FA) (buffer A) and CH3CN/FA 0.1% (80/20, buffer B). Tryptic peptides were eluted at a flow rate of 300 nL/min using a three-step linear gradient: from 2 to 40% B over 123 min, from 40 to 100% B in 1 and 10 min at 100% B. The mass spectrometer was operated in positive ionization mode with capillary voltage and source temperature set at 1.9 kV and 275 °C, respectively. The samples were analyzed using HCD (higher-energy collision dissociation) method. The first scan (MS spectra) was recorded in the Orbitrap analyzer (R = 120,000) with the mass range m/z 400–1800. Then, 20 scans were recorded for MS2 experiments. Singly charged species were excluded for MS2 experiments. Dynamic exclusion of already fragmented precursor ions was applied for 30 s, and an exclusion mass width of ± 10 ppm. Peptide isolation was achieved in the quadrupole with an isolation window of 1.6 m/z. Fragmentation occurred with a HCD collision energy of 28%. Daughter ions were analyzed in the Orbitrap with a resolution of 15,000. The maximum injection times were 30 ms and 22 ms for MS and MS2 analyses, respectively. All measurements in the Orbitrap analyzer were performed with on-the-fly internal recalibration (lock mass) at m/z 445.12002 (polydimethylcyclosiloxane).

### Protein Quantification

Raw data were imported in Progenesis LC-MS/MS software (Waters, ver. 4.1, UK). A 2-dimensional map was generated for each sample (retention time versus *m/z* ratio). The spots present on the 2D maps were then aligned. Monocharged ions and those with a charge greater than 5 were excluded from the analysis. MS/MS spectra from selected peptides were exported for peptide identification with Mascot (Matrix Science, ver. 2.2.04) against the *S. aureus* NCTC 8325 database available online (https://www.ncbi.nlm.nih.gov/datasets/taxonomy/93061/). Database searches were performed with the following parameters: 1 missed trypsin cleavage site allowed; variable modifications: carbamidomethylation of cysteine and oxidation of methionine. Mass tolerances for precursor and fragment ions were both set at 5 ppm. Then, Mascot search results were imported into Progenesis. Peptides with an identification score greater than 13 are retained. Proteins identified with less than 2 peptides were discarded. For each condition (Wild type and mutants), the total cumulative abundance of the protein was calculated by summing the peptides abundances. The software compared the intensity of the isotopic mass of all the ions for each condition. The abundances were normalized to perform a relative quantification of each protein between the conditions. The MS proteomics data have been deposited to the ProteomeXchange Consortium via the PRIDE partner repository with the date set identifier PXD060402.

### Statistical analyses of proteomic data

Significance analyses of differentially regulated proteins were performed with Perseus software (version 2.0.10.0.). Proteins normalized abundance (from Progenesis) were transformed to log10 space. Significantly regulated proteins between the different strains (Wild type, Mutant and complemented) were identified by two sample t test with a *p-value* ≤ 0.05 and volcano plot was generated.

### Functional annotation and functional enrichment analysis

The proteins were functionally annotated via NCBI, UniProt and Kyoto Encyclopedia of Genes and Genomes (KEGG) functional resources and added to the Perseus software. The significantly up- and down-regulated proteins from the three different comparisons were tested for functional enrichment based on KEGG pathways by using the Fisher’s exact test. Annotation terms were considered significant on the basis of an enrichment factor ≥ 1, an intersection size > 2 and a p-value ≤ 0.2.

### RNA extractions, Northern blots, RNA half-life and RT-qPCR

The cells were collected and pelleted for 10 min at 4°C, at 4500 rpm, then resuspended in RNA lysis buffer consisting of 0.5% SDS, 20 mM sodium acetate, 1 mM EDTA, pH 5.5. Total RNA extraction was performed as previously described (27). Northern blot assays were conducted following established protocols (27). Membranes were hybridized with specific 32P-labeled probes (Supplementary Table S2) in ExpressHyb solution (Ozyme) and washed according to recommendations from manufacture. The membranes were exposed and scanned with Typhoon FLA 9500 scanner (GE Healthcare). Image quantification was carried out using ImageQuant Tool 7.0. To determine the half-life of sRNA, rifampicin (200 µg/ml) was employed as the standard treatment to halt transcription. Approximately 4 ml of each strain was collected prior to and at 2, 5, 10, 20, 30, 40, and 60 minutes after rifampicin addition. These samples were centrifuged, and the pellets were frozen in liquid nitrogen and subsequently stored at −80°C for total RNA extraction. For quantitative real-time PCR (qRT-PCR), the total RNA extraction samples were treated with the DNase I Amplification Grade Kit (Invitrogen) to remove any residual DNA. cDNA preparations and qRT-PCR experiments were conducted as previously described (23), with the *gyrB* gene serving as the normalization control.

### *In vitro* transcription, RNA labeling and gel retardation assays

Gel-shift assays were performed as described in (23). RNAs were synthesized from PCR-generated DNA using MEGAscript T7 kit (Ambion). The transcription template was amplified from HG003 genomic DNA and forward primers included T7 promoter sequences (Supplementary Table S2). Then, RNAs were labeled at 5′-end using [γ-32P] ATP (Amersham Biosciences) and T4 polynucleotide kinase (Invitrogen). Both labeled and unlabeled RNAs were purified on a 5% acrylamide-urea gel, eluted in elution buffer (20 mM Tris-HCl, pH 7.5; 250 mM NaCl; 1 mM EDTA; 1% SDS) at 37°C. The RNA was then ethanol-precipitated, quantified using a Qubit fluorometer (Thermo Fisher Scientific), and stored at −80°C. Gel-shift assays were conducted as outlined in (23). The RNAs were denatured in a solution of 50 mM Tris/HEPES (pH 7–7.5) and 50 mM NaCl for 2 minutes at 80°C, followed by refolding for 10 minutes at 25°C upon the addition of MgClLJ to a final concentration of 5 mM. Reactions were carried out in a buffer containing 50 mM Tris-HCl (pH 7.5), 50 mM NaCl, and 5 mM MgClLJ for 20 minutes at 25°C. Approximately 0.05 pmole of labeled RsaOI or RsaOI-del were incubated with varying concentrations of *lacD* mRNA. The samples were supplemented with 10% glycerol and loaded onto a native 4% polyacrylamide gel containing 5% glycerol. The gels were subsequently dried and visualized using a Typhoon FLA 9500 scanner (GE Healthcare).

## RESULTS

### *rsaOI* expression is triggered by low pH, improving survival in acidic stress conditions

To identify sRNAs potentially involved in adaptation to a low pH environment, we performed RNA-seq analysis, comparing the transcriptome profiles of the HG003 strain under neutral (pH 7) and acidic (pH 4) conditions. For this purpose, total RNA was extracted from two HG003 cultures in the exponential growth phase, either maintained at neutral pH or exposed to acidic stress for 30 minutes, as outlined in the Materials and Methods.

Several genes identified as part of the acid stress response regulon showed increased expression (Supplementary Table 3). For instance, the *ure* operon encoding urease subunits A, B and C (*ureA*, *ureB* and *ureC*), which plays a role in maintaining intracellular pH, was upregulated (13). Urease activity is known to be the major acid-resistance mechanism (2). Additionally, the *kdpDE* operon, which encodes a two-component system involved in both acid and osmoprotection, was stimulated (28). Genes associated with histidine import and metabolism, essential for growth at low pH, were also upregulated (8). In agreement with previous studies on sudden inorganic stress, we observed a general reduction in the expression of ribosomal protein genes, and pyrimidine ribonucleotide biosynthesis pathway, which likely reflects the lowered growth rate after acidification (12). Among the genes affected by low pH, *rsaOI* showed one of the most significant responses, with acidic conditions leading to more than a 14-fold accumulation of RsaOI (Figure 1A and Supplementary Table 4). RsaOI is approximately 250 nucleotides (NT) long and represents the longest sRNA that we previously identified in strain N315 (29). Using RACE (rapid <u>a</u>mplification of <u>c</u>DNA <u>e</u>nds), we mapped the 5’-end of *rsaOI* from HG003 strain to position 565901 in the NCTC8325 genome, which corresponds to the location in N315 (29). The *rsaOI* gene is present in all analyzed *S. aureus* strains, exhibiting 100% sequence conservation across isolates (Supplementary Figure 1). Moreover, this sRNA is also found in other staphylococcal species, including *Staphylococcus schweitzeri* and *Staphylococcus argenteus* with a variable sequence conservation (98,4 to 95,7% sequence identity respectively) (Supplementary Figure 1). In *S. aureus* and related staphylococcal species, compensatory mutations confirm that the last 30 nucleotides of RsaOI form a hairpin structure, likely functioning as a rho-independent transcription terminator. This is consistent with results showing that *rsaOI* transcription is not affected by *rho* deletion (30). Moreover, in HG003 strain, *rsaOI* has its own promoter and transcription terminator, with no open reading frame on the opposite strand. This sRNA can therefore be classified as a *bona fide* sRNA (31). *rsaOI* gene is located between *proP* gene, encoding for an osmolyte transporter that facilitates bacterial adaptation to osmotic stress (32) and operon *vraABC* involved in fatty acid metabolism, and shown to be-upregulated in staphylococcal strains with increased resistance to vancomycin (33) (Figure 1B).

**Figure 1:**
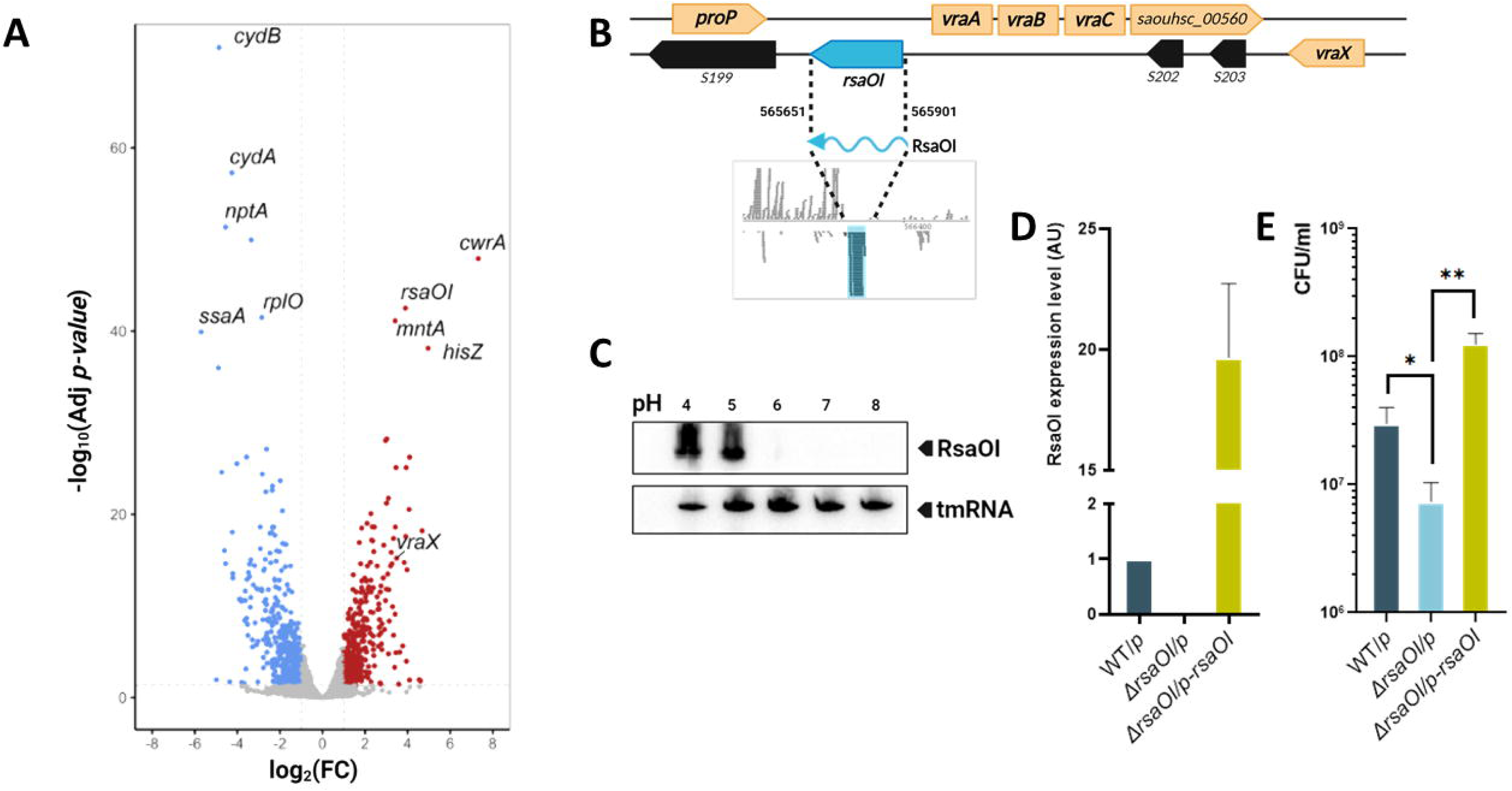
Induced by low pH, *rsaOI* contributes to survival under acidic stress. (**A**) Visualization of the most significantly accumulated mRNAs under acidic (pH 4) and neutral (pH 7) conditions using a Volcano plot. The genes with the highest differential expression are highlighted and labeled. (**B**) Genomic localization of *rsaOI* gene in HG003 strain, situated between *proP* and *vraABC* operon. (**C**) RsaOI expression in response to pH range was analyzed by Northern blot analysis using *rsaOI* specific probe, with tmRNA used as loading control. (**D**) *rsaOI* expression in HG003 WT, Δ*rsaOI* (both carrying pICS3 plasmid; WT*/p* and Δ*rsaOI/p*) or complemented strain (Δ*rsaOI*, carrying pICS3-*rsaOI* plasmid) was analyzed by qPCR, normalized to the control gene *gyrB*, and calculated using 2^−ΔΔCt^ method for relative quantification. (**E**) Strains from pannel (**D**) were cultured in LB medium for 2 hours, then pelleted and resuspended in LB buffered to pH5. After 2 hours at pH5, samples were collected and plated to determine CFU counts. Statistical analysis was conducted using Student’s t-test. Error bars represent the average of three independent experiments. Statistical significance is indicated by bars and asterisks as follows: *, P<0.01; **, P<0.05.

To validate the results obtained from the RNA-seq analysis of the effect of pH on *rsaOI* expression, Northern blot and RT-qPCR analyses were performed. They both showed that the increase in RsaOI levels correlated with a decrease in the pH of medium: the highest accumulation of RsaOI was observed at pH 5 and 4, corresponding to 200-fold and 400-fold increases, respectively, compared to RsaOI levels at pH 7 (Figure 1C and Supplementary Figure 2). This supports the low pH-dependent expression of *rsaOI*.

We monitored the *rsaOI* expression profile during growth and found that *rsaOI* expression increased significantly during the stationary growth phase in nutrient-rich media in HG003. This pattern was conserved across other tested strains (N315, Newman, and USA300; Supplementary Figure 3A and 3B), these data align with the highest expression of *rsaOI* at stationary phase reported for *rsaOI* from N315 strain grown in nutrient-rich media BHI (29). We hypothesised that RsaOI increased levels was due to acidification of the medium as a result of glucose consumption during bacterial growth (5). Consequently, *rsaOI* expression was analyzed in LB medium with or without glucose, and under buffered pH conditions preventing acidification of the medium. Growth of *S. aureus* in LB medium with glucose led to a progressive increase in RsaOI levels, which was prevented by buffering the medium to neutral pH (Supplementary Figures 4A and 4B). *rsaOI* expression was also assessed under stress conditions encountered by bacteria during infection. Among the conditions tested, only the acidic stress significantly induced *rsaOI* expression, either by inorganic (HCl) or organic (acetic acid) acids (Supplementary Figure 5). These findings support that *rsaOI* expression is induced by low pH, regardless of the origin of pH lowering.

We further investigated the role of RsaOI in *S. aureus* acid resistance. To assess this, HG003 wild-type (WT) and *rsaOI* deleted (Δ*rsaOI*) strains were exposed to acid stress at pH5, followed by plating to measure viability. Additionally, the *rsaOI* deletion was complemented with high copy plasmid pICS3-*rsaOI,* expressing a wild-type copy of the *rsaOI* gene under the control of its native promoter (Δ*rsaOI/p-rsaOI*) (Figure 1D). Colony-forming unit (CFU) counts revealed that *rsaOI* expression significantly improved survival rates compared to the Δ*rsaOI* strain (Figure 1E).

### *rsaOI* expression is affected by envelope stress and regulated by the VraSR two-component system

To explore the mechanism underlying RsaOI accumulation, we tested whether the increase in *rsaOI* expression under acidic conditions was due to its promoter activation. For this, we constructed a plasmid with GFP under control of *rsaOI* promoter (pCN38-P*rsaOI*-*gfp*). As a control, a plasmid with GFP regulated by the *tufA* promoter was employed (pCN38-P*tufA*-*gfp*). Acidic conditions did not affect *tufA* promoter activity but enhanced the fluorescence associated with the *rsaOI* promoter (Figure 2A). To further validate this finding, we created a plasmid with the *rsaOI* expression driven by the constitutive promoter *amiA* (pICS3-*PamiA-rsaOI*). Expectedly, a decrease in pH did not impact RsaOI levels when *rsaOI* was under the control of the constitutive promoter, whereas there was an induction of *rsaOI* expression when regulated by its native promoter (pICS3-*rsaOI*) (Supplementary Figure 6A). Additionally, we found that the decrease in pH did not affect RsaOI stability (Supplementary Figure 6B). Taken together, these results indicate that the increase in RsaOI levels under acidic conditions is driven transcriptionally by enhanced *rsaOI* promoter activity.

**Figure 2:**
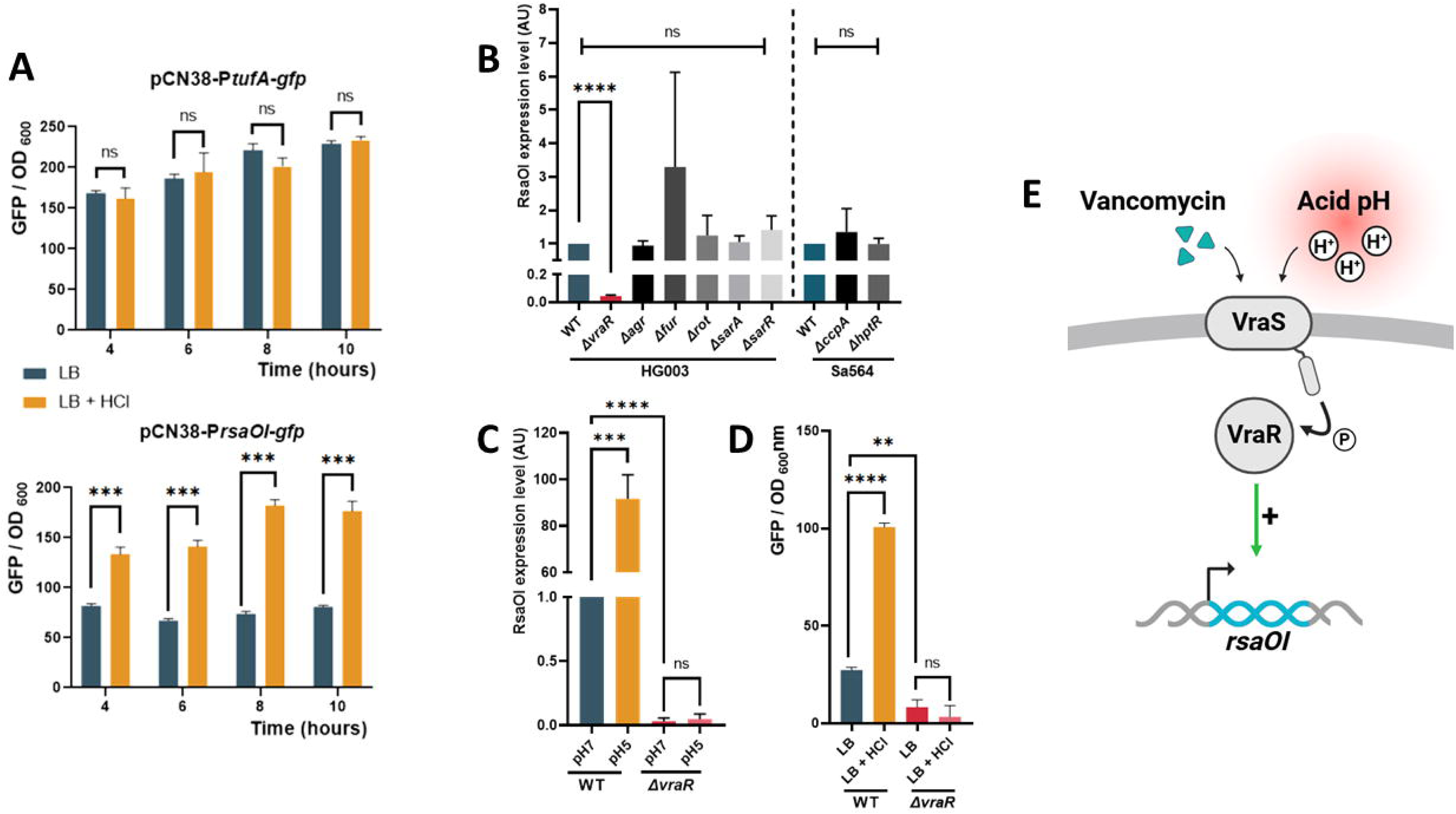
VraR affects *rsaOI* expression. (**A**) Fluorescence levels of GFP under control of either *tufA* or *rsaOI* promoters. Cells were grown for 2 hours in LB medium, followed by addition of HCl. All statistical analyses were performed using Student’s t-test. Error bars represent the average of three independent experiments. Statistical significance is indicated by bars and asterisks as follows: ***, P<0.001. (**B**) Analysis of *rsaOI* levels in HG003 and Sa564 wild-type strains (WT) and their isogenic mutants. Relative quantification of *rsaOI* expression levels were measured by qPCR, normalized to the control gene *gyrB*, and calculated using 2^−ΔΔCt^ method. (**C**) *rsaOI* expression in WT and in Δ*vraR* mutant according to pH. Cells were cultured in LB medium for 2 hours, then pelleted and resuspended in LB buffered with 100 mM HEPES at pH 5 or pH 7 for 30 minutes. (**D**) Fluorescence levels of GFP under control of *rsaOI* promoter in both HG003 WT and Δ*vraR* strains. Cells were grown for 2 hours in LB medium, followed by HCl addition. Statistical analysis was performed using Student’s t-test. Error bars represent the average of three independent experiments. Statistical significance is indicated by bars and asterisks as follows: ***, P<0.001; ****, P<0.0001. (**E**) Illustration of the induction mediated by the TCS VraSR on the expression of the sRNA *rsaOI* during acid stress or in the presence of the glycopeptide antibiotic vancomycin.

Interestingly, previous studies have reported that *rsaOI* expression was also increased in the presence of vancomycin (34, 35), but the regulatory network involved was not identified, although involvement of VraSR system was supposed (34). Therefore, to identify the factor responsible for RsaOI accumulation under acidic conditions, we tested the *vraR* mutant together with several mutants lacking transcription regulators and TCSs for *rsaOI* expression. All tested strains except for the *vraR* mutant, showed elevated *rsaOI* expression under acidic condition (Figure 2B). The absence of VraR prevented the accumulation of RsaOI whether the pH of the medium was 5 or 7, indicating that VraR is required for *rsaOI* expression (Figure 2C). Similarly, a *vraS* mutant yielded the same results, further supporting the role of this TCS in *rsaOI* regulation (data not shown). VraS functions as a histidine kinase, while VraR is the response regulator of the VraSR (VraR/VraS) TCS that governs the staphylococcal response to perturbation in cell wall synthesis (36). This TCS was initially identified for its role in vancomycin resistance (37).

To test whether acidic stress activates the VraSR system, we evaluated the expression of *vraX* and *cwrA* genes, which are directly regulated by VraSR (38, 39). In accordance with transcriptomic data (Figure 1A), the expression of both *vraX* and *cwrA* was upregulated under acidic stress in wild-type HG003 strain (Supplementary Figure 7A and 7B). However, the expression of both genes was strongly downregulated in a *vraR* mutant (Supplementary Figure 7A and 7B). Those data indicate that *cwrA* and *vraX* are part of an acid stress regulon directly regulated by VraSR. Additionally, we observed that acidic conditions slightly led to higher *vraSR* mRNA levels, consistent with reported findings (12) (Supplementary Figure 7C).

To confirm that the *rsaOI* regulation by VraSR occurs *via* its promoter, HG003 wild-type strain and Δ*vraR* mutant were transformed by plasmid containing GFP reporter driven by the *rsaOI* promoter (pCN38-P*rsaOI*-*gfp*). No induction of fluorescence under acidic condition was observed in Δ*vraR* mutant in contrast to the wild type strain promoter (Figure 2D). Altogether, our results demonstrate that in addition to responding to cell wall stress caused by previously reported cell wall-damaging agents (40), VraSR TCS mediates response to low acidic growth conditions (Figure 2E) and induces the transcription of *rsaOI* in response to a decrease in external pH.

### RsaOI impacts *S. aureus* susceptibility to glycopeptide antibiotics

Since *rsaOI* was identified as part of the regulon of VraSR, a TCS involved in response to cell wall stress and in resistance to cell wall active-antibiotics, we investigated whether RsaOI affected *S. aureus* susceptibility to glycopeptide antibiotics. For this, we compared the *rsaOI* mutant strain and its parental wild-type strain HG003 for susceptibility to vancomycin. Additionally, *rsaOI* deletion was complemented with a plasmid pICS3-*rsaOI*. While no differences in growth were observed between the strains in the absence of vancomycin, the loss of *rsaOI* led to improved growth in the presence of the vancomycin, compared to the wild-type strain (Figure 3A and 3B). Complementation of the *rsaOI* deletion restored the reduced growth in the presence of vancomycin, confirming that RsaOI contributes to *S. aureus* susceptibility to this antibiotic. Similar results were obtained using a spot population analysis profile assay (Supplementary Figure 8) (41).

**Figure 3:**
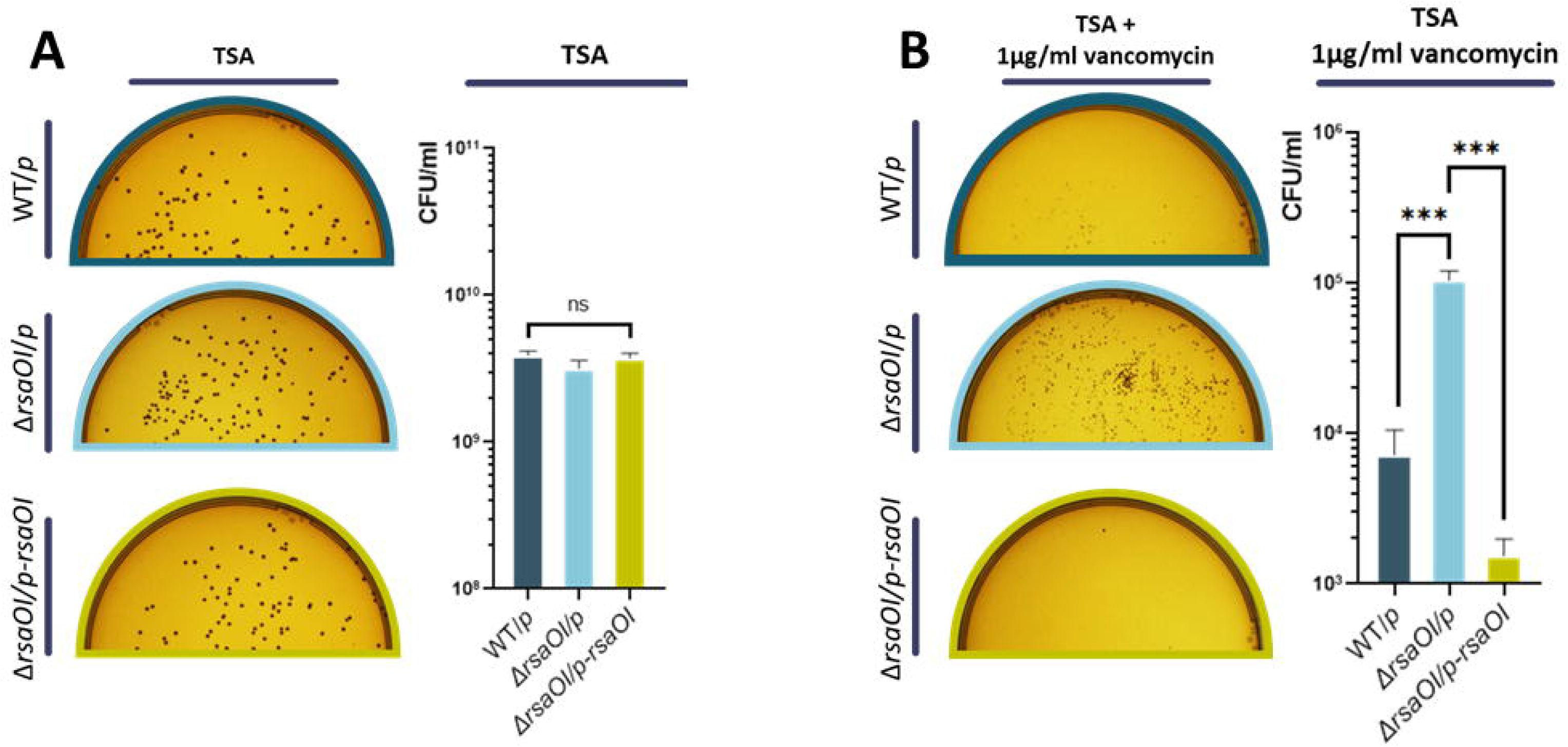
RsaOI modulates *S. aureus* resistance to vancomycin. Ten-fold serial dilutions of overnight cultures of WT HG003, Δ*rsaOI* or complemented strain (Δ*rsaOI/p-rsaOI*) were plated and incubated for 24 hours at 37°C. The 10^-6^ dilution are shown for TSA condition (**A**), and the 10^-4^ dilution for TSA supplemented with 1µg/ml vancomycin (**B**). Diagrams show colony growth quantification. Statistical analysis was performed using Student’s t-test. Error bars represent the average of three independent experiments. Statistical significance is indicated by bars and asterisks as follows: ***, P<0.001.

### Identification of protein levels affected by RsaOI

Altogether, these RsaOI features prompted us to investigate its targets and regulatory mechanism. To identify RsaOI-dependent changes in the abundance of individual proteins, we analyzed by quantitative proteomics which protein levels were affected by RsaOI expression. For this purpose, we compared the proteome of HG003 wild-type (WT), Δ*rsaOI* containing an empty vector (WT/p vs Δ*rsaOI*/p, respectively) and Δ*rsaOI* complemented with the pICS3-*rsaOI* vector (Δ*rsaOI*/*p-rsaOI*). To ensure *rsaOI* expression from its native promoter in the HG003 WT and complemented strains, bacteria were grown under low pH conditions (Supplementary Figure 9A). From 2,573 annotated putative proteins in *S. aureus* NCTC 8325 strain (42), 1,112 proteins were identified in HG003 wild-type, 1,012 in Δ*rsaOI* and 996 in *rsaOI* complemented strain. Tryptic peptides were analysed by nLC-MS/MS (n = 4 per condition). We performed a functional enrichment analysis based on the significantly up- or down-regulated proteins in comparison Δ*rsaOI*/p vs WT/p, and Δ*rsaOI*/p vs Δ*rsaOI*/*p-rsaOI*. Overall, carbon metabolism proteins were upregulated in the Δ*rsaOI* strain compared to the WT or *rsaOI* complemented strains (Supplementary Figure 9B and 9C). For example, among these proteins, we found Fda (Fructose-1,6-bisphosphate aldolase) involved in glycolysis, SucA (2-oxoglutarate dehydrogenase E1 component) in the TCA cycle, as well as LacD (Tagatose 1,6-diphosphate aldolase) part of galactose metabolism. This finding could be relevant in light of the impact of metabolism on medium acidification.

Focusing on proteins with differential abundance between deletion strain Δ*rsaOI*/p and complemented strain Δ*rsaOI*/p-*rsaOI* using *t*-test (*p-value* ≤0.05), we highlight that a total of 129 proteins were significantly differentially regulated, including 30 proteins with a fold change greater than 2, (21 up and 9 downregulated in Δ*rsaOI* strain; Figure 4A and Supplementary Table 5). Interestingly, the amount of Atl, which was recently shown to be repressed by RsaOI (34), was found to be increased in Δ*rsaOI* strain. Notably, the transcriptional regulator of RsaOI, VraR, was found to be less abundant in the strain expressing *rsaOI*. The inhibition of VraR protein levels is particularly noteworthy given the significantly reduced survival of a strain expressing *rsaOI* in the presence of glycopeptides. Altogether, while the proteomic analysis does not elucidate the underlying regulatory mechanisms, it underscored a substantial impact of RsaOI on the levels of numerous proteins under acidic conditions.

**Figure 4:**
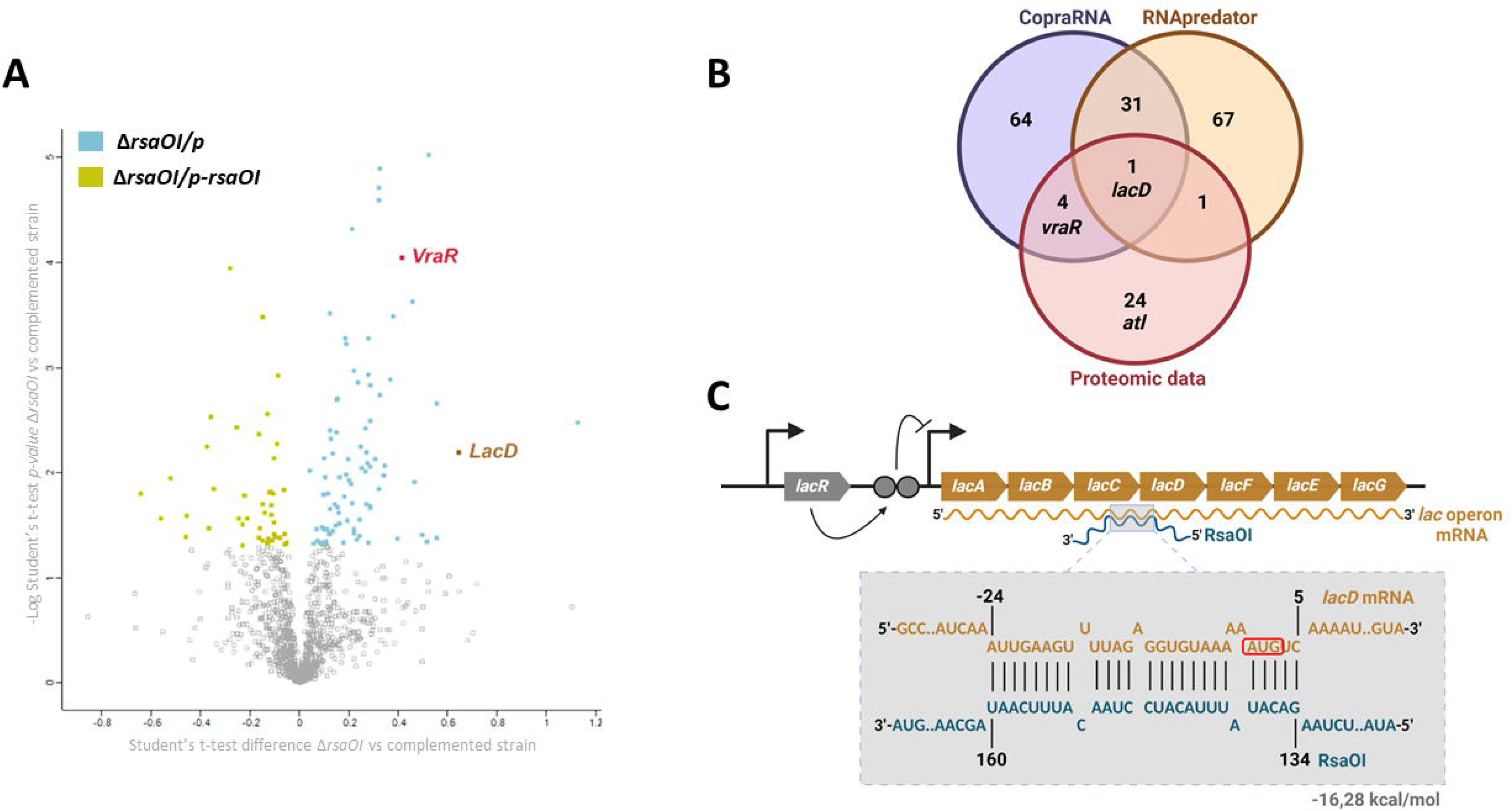
Investigation of the potential direct targets of RsaOI based on *in vivo* proteomic data and *in silico* computational approaches. (**A**) Volcano-plot showing protein relative abundances between the Δ*rsaOI* strain containing a pICS3 vector (Δ*rsaOI/p)* and the complemented strain (Δ*rsaOI/p-*Δ*rsaOI*) in relation to the *t*-test *p*-value (<0.05). VraR and LacD are significantly more abundant in Δ*rsaOI* strain. (**B**) Venn diagram illustrating the overlap between experimentally identified and *in silico* predicted RsaOI target candidates. Experimentally identified candidates, representing proteins with significantly different abundances between *rsaOI* mutant strain and complemented, are highlighted in red. The top 100 predicted targets from CopraRNA (43) are shown in blue, while the top 100 predicted by RNA predator (44) are depicted in yellow. Shared targets are represented in the overlapping regions. (**C**) Organization of the *lac* operon in the HG003 strain and the predicted base-pairing interaction between RsaOI and *lac* operon mRNA, generated using IntraRNA software (78). The start codon of *lacD* is highlighted in red, and position +1 corresponds to the transcription start site for RsaOI.

### Integration of experimental data and computational prediction highlights potential direct targets of RsaOI

The effect of sRNAs on the protein abundance can occur through sRNA-pairings with their corresponding mRNA or indirectly through the activity of sRNAs on regulatory proteins. To identify potential direct targets of RsaOI among candidates identified by proteomic approach, an *in silico* analysis was conducted using CopraRNA and RNA predator, which predict sRNA-mRNA interactions (43, 44). These software tools generated a list of putative sRNA targets based on the likelihood of direct interaction with RsaOI through sequence complementarity, while also considering the secondary structure of both interacting partners. We focused on predicted targets that overlapped between the *in-silico* predictions and our proteomic data. Notably, we identified 20 mRNA targets: 10 predicted by CopraRNA and 10 by RNApredator (Figure 4B and Supplementary Figure 10A). Interestingly, among these potential targets, *vraSR* mRNA, encoding the transcriptional regulator of *rsaOI*, was predicted to interact with RsaOI by CopraRNA (Figure 4B). The predicted interaction is located in *vraS* coding sequence, 87 nucleotides upstream of translational initiation site of *vraR* (Supplementary Figure 10B).

Regarding the potential regulation of VraSR by RsaOI, we hypothesized that differences in glycopeptide survival could result from RsaOI-mediated inhibition of VraR level. To test this, a vancomycin survival assay using *vraR* deletion strains was conducted, with or without RsaOI expression driven by the constitutive *amiA* promoter (Supplementary Figure 11). The deletion of *vraR* significantly increased bacterial susceptibility to vancomycin, as previously described. Notably, *rsaOI* expression in a *vraR* deletion mutant further affected survival, indicating that RsaOI influences vancomycin tolerance through the regulation of an additional target.

The Venn diagram analysis revealed only a single hit between the two bioinformatics tools and the proteomics data: *lacD* (SAOUHSC_02452) (Figure 4B and 4C). The *lacD* gene is within the seven-gene *lac* operon (*lacABCDFEG*) involved in galactose metabolism through the tagatose pathway. RsaOI is predicted to pair with 25 nucleotides of the *lacD* mRNA, which includes its ribosome binding site (RBS) and start codon (Figure 4C). It should be noted that this region is conserved in other staphylococcal species, such as *S. argentus* and *S. schweitzeri*, where *rsaOI* is also present (Supplementary Figure 12). Altogether, the compilation of *in vivo* data showing that of RsaOI reduces the LacD protein levels and *in silico* predictions of RsaOI binding at the *lacD* RBS, suggested a post-transcriptional mechanism of *lacD* regulation involving RsaOI.

### RsaOI controls expression of *lacD* at translational level

To confirm the *in silico* prediction of RsaOI binding the *lac* mRNA in the RBS region of *lacD*, the formation of the RsaOI-*lacD* mRNA duplex was evaluated by electrophoretic mobility shift assay (EMSA). A 250-nt-long mRNA fragment of *lac* mRNA containing the predicted RsaOI binding site was produced. RsaOI was able to form a complex with the *lacD* mRNA fragment (Figure 5A), with 50% of complex at 50 pmol of *lacD* mRNA (Figure 5B). The specificity of this interaction was confirmed by the lack of binding interference from an excess of yeast tRNA (Figure 5A). We constructed a mutant version of RsaOI (RsaOI-mut), which lacks 30 nucleotides predicted to be involved in the interaction site with *lacD* mRNA. RsaOI-mut was unable to form a duplex with *lacD* mRNA, indicating that these nucleotides are required for direct interaction (Figure 5A and 5B).

**Figure 5:**
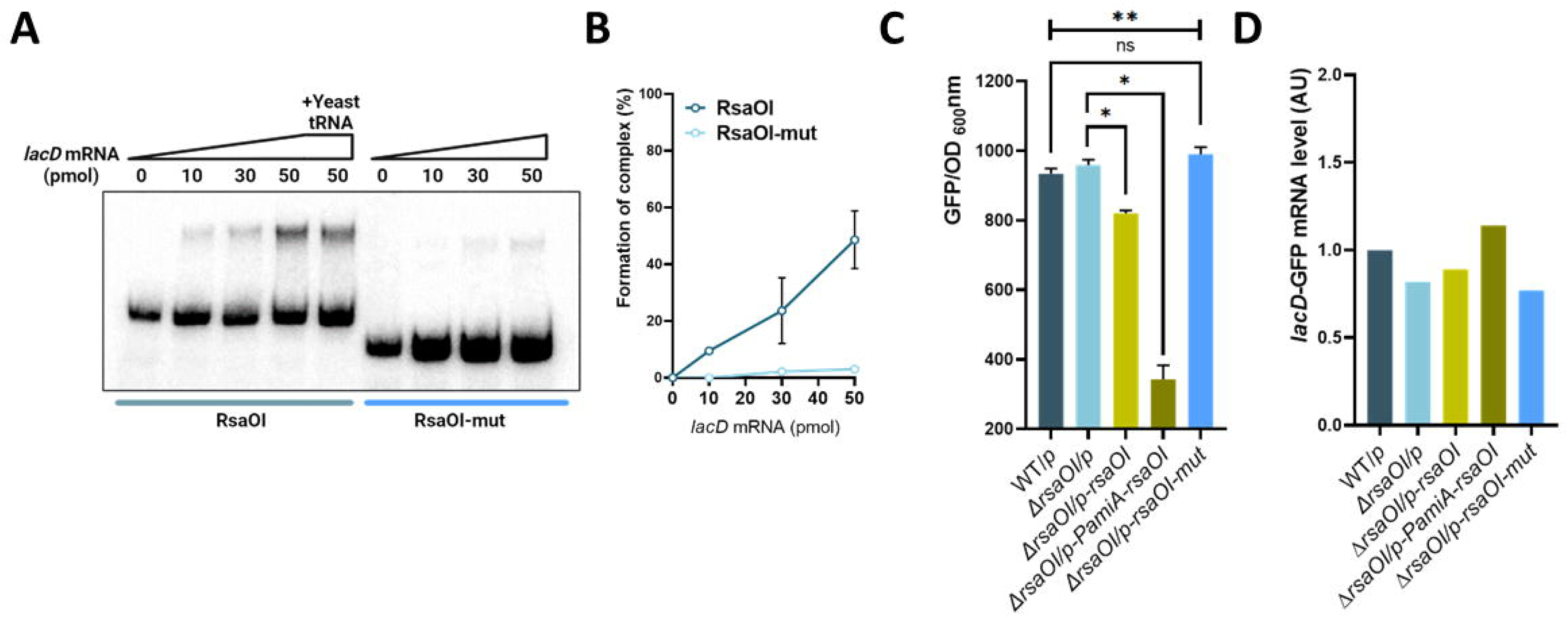
RsaOI inhibits *lacD* expression. (**A**) Complex formation between RsaOI and *lacD* mRNA was analyzed by native gel retardation assays. Gel shift assays of purified, labeled RsaOI and RsaOI-mut were performed with increasing concentrations of *lacD* mRNA. (**B**) Quantification of complex formation from (A) was performed by ImageQuant Tools 7.0. (**C**) The effect of RsaOI on *lacD* expression was examined using *gfp* gene reporter assay. *S. aureus* WT and Δ*rsaOI* strains containing the pCN33*-PtufA-lacD-gfp* fusion plasmid were co-transformed with pICS3, pICS3 expressing *rsaOI* or *rsaOI-mut* under control of its endogenous promoter, or *rsaOI* under constitutive promoter *amiA*. The fluorescent intensity was measured after 8 hours of growth. Statistical analysis was conducted using a Kruskal-Wallis non-parametric test across all conditions followed by Student’s t-test to determine significant differences among conditions. Error bars represent the average of four independent experiments. Statistical significance is indicated by bars and asterisks as follows: *, P<0.05; **, P<0.01. (**D**) In parallele with the panel (C) experiment, strains were collected and pelleted, and relative quantification of *lacD-gfp* fusion mRNA expression levels were measured by qPCR, normalized to the control gene *gyrB*, and calculated using 2^−ΔΔCt^ method.

Next, the effect of RsaOI binding to the *lacD* RBS on *lacD* regulation was evaluated *in vivo*. For this, the sequence of *lacD* mRNA used for *in vitro* study of interaction was fused in-frame with GFP reporter and positioned under the control of the constitutive *tufA* promoter to bypass the regulation on transcriptional level. Plasmid expressing *lacD*-GFP reporter fusion was introduced in both the HG003 strain and its isogenic derivative Δ*rsaOI*. *rsaOI* was expressed under control of its native promoter (pICS3-*rsaOI*) or a constitutive promoter (pICS3-*PamiA-rsaOI*). The effects of RsaOI on *lacD* expression were evaluated by measuring fluorescence intensity through quantitative microplate assays. To permit *rsaOI* expression under control of its native promoter, we cultured the cells in BHI medium, which allows acidification of the medium (Supplementary Figure 13). Expression of *rsaOI* reduced the fluorescence produced by *S. aureus* cells containing *lacD-*GFP translational fusion (Figure 5C). The decrease of fluorescence was correlated with *rsaOI* expression level, since the high level of *rsaOI* expressed from a strong *amiA* promoter resulted in more pronounced fluorescent inhibition. Moreover, the expression of *rsaOI*-mut allele (Supplementary Figure 13) unable for *lacD* mRNA binding *in vitro* (p-*rsaOI-mut*) had no effect over the fluorescence, thereby confirming the specificity of the regulation. To note, that for unknown reasons, we were unable to introduce the vector expressing the *rsaOI*-mut under the control of the constitutive *amiA* promoter in *S. aureus*. To determine whether RsaOI affected *lacD* by altering its mRNA levels, we measured the amount of *lacD-gfp* mRNA depending on RsaOI expression (Figure 5D). No significant effect of RsaOI on mRNA levels was noticed suggesting that RsaOI acts through an antisense mechanism, leading to translational repression of *lacD*.

### RsaOI impairs galactose utilization in *S. aureus*

Galactose is usually metabolized via the well-known Leloir pathway (45), where D-galactose is converted through D-galactose-l-phosphate and D-glucose-l-phosphate to D-glucose-6-phosphate. However, *S. aureus* lacks the enzymes of the Leloir pathway and in this bacterium as opposed to other staphylococcal species such as *S. intermedius, S. saprophyticus* and *S. xylosus*, galactose is converted to D-galactose 6-phosphate, which is further metabolized through tagatose derivatives (D-galactose-6-phosphate -> D-tagatose-6-phosphate -> D-tagatose-1,6-diphosphate -> D-glyceraldehyde-3- phosphate + dihydroxyacetone-phosphate) (46) (Figure 6A). Thus, *S. aureus* metabolize galactose exclusively *via* tagatose pathway (47), which is mediated by the *lac* operon (Figures 4 and 6A).

**Figure 6:**
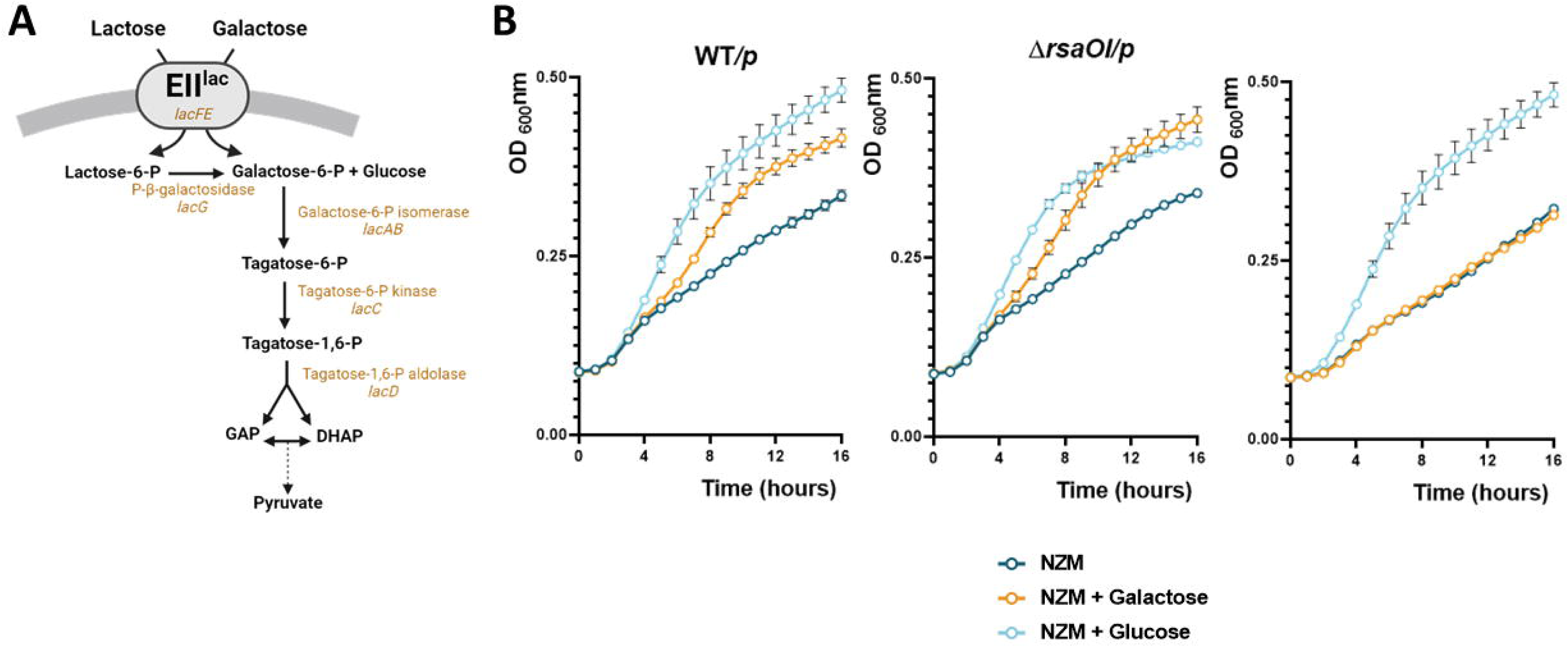
*rsaOI* overexpression impairs using of galactose as an external source of carbons. (**A**) Illustration of the tagatose pathway, a metabolic pathway that catabolizes galactose or lactose through the *lac* operon. (**B**) Growth curves of *S. aureus* HG003 WT, Δ*rsaOI*, or complemented strain expressing *rsaOI* under constitutive promoter *amiA* were cultured in NZM media supplemented with glucose or galactose at 11mM. Growth was followed during 16 hours using a Biotek microplate reader. Error bars represent the average of three independent experiments.

Since, the tagatose pathway is the sole route for galactose utilization in *S. aureus* and regarding the effect of RsaOI on *lacD* expression, we evaluated an impact of RsaOI on bacterial growth in medium supplemented with galactose. We used NZM medium, which lacks external sugar, and NZM supplemented with either glucose or galactose (Figure 6B). In these media, we compared the growth of the *S. aureus* HG003 WT, Δ*rsaOI*, and a strain that constitutively expresses *rsaOI* in a condition-independent manner (Δ*rsaOI/p-PamiA-rsaOI*). All strains showed similar growth in NZM alone. Additionally, the supplementation with glucose significantly enhanced growth of all strains. However, when galactose was added, we observed differences in growth depending on *rsaOI*. Indeed, galactose improved the growth of both WT and Δ*rsaOI* strains. In contrast, the strain constitutively expressing RsaOI exhibited no growth improvement with galactose compared to NZM alone (Figure 6B). This suggests that expression of RsaOI inhibits galactose utilization likely due to the repression of the *lac* operon.

## DISCUSSION

Acid stress is a common challenge encountered by *S. aureus* throughout its life cycle. This stress can be either endogenous, triggered by sugar metabolism, or exogenous, such as exposure to host defences or antimicrobials. To counteract the deleterious effect of pH lowering, *S. aureus* utilizes specific mechanisms and alters its transcriptome by upregulating the acid resistance regulon including genes involved in urea degradation, amino acid uptake, or factors that modify the membrane charge (2).

In this study, we examined gene expression changes in response to acid stress. In addition to identifying genes belonging to the characteristic acid stress response regulon (2), transcriptomic analysis revealed a small RNA, RsaOI that accumulates significantly under acidic pH conditions. Further analysis showed that the expression of *rsaOI* is induced by low pH, regardless of either inorganic or organic acid is used and whether the acidification is caused by external acid or results from glucose metabolisms. We identified VraR, the response regulator of the VraSR two-component system, as a key activator of *rsaOI* expression under acidic conditions (Figure 7). Specifically, VraR activates the *rsaOI* promoter in response to pH decrease. VraSR is homologous to the LiaSR TCSs found in other Gram-positive bacteria, such as *B. subtilis*, *Streptococcus mutans*, and *Enterococcus faecalis* (48, 49). In accordance with TCS from *B. subtilis* and *L. lactis* and other Gram-positive bacteria, the staphylococcal system also contains a third component (VraT) corresponding to LiaF (50). In response to an unknown signal, likely induced by exposure to cell wall-targeting agents, VraS undergoes autophosphorylation, leading to its activation and subsequent phosphorylation of VraR (40, 51). The phosphorylated VraR dimerizes and binds to the promoter of its own operon (*vraUTSR*) as well as the promoters of numerous genes that constitute the so-called cell wall stress stimulon (CWSS)(52)(Figure 7). Genes induced by this mechanism are involved in diverse cellular processes, including DNA replication and repair, carbohydrate metabolism, and, most notably, a substantial number of genes essential for cell wall biosynthesis and remodeling. This regulation enables an adaptive response to maintain cell envelope integrity under stress conditions (51). However, both the stimuli that activates VraSR and the specific configuration of VraSR-regulated genes can vary significantly, depending on the strains and experimental procedures used (51, 53)(Figure 7). While, in other Gram-positive bacteria, the *liaSR* homologs have been shown to be involved in acid stress response (50), the VraSR system of *S. aureus* was initially reported to be induced by cell wall-targeting antibiotics, including vancomycin, teicoplanin and β-lactams, but not by general stresses, such as acid pH (40, 51). However, our transcriptome and follow-up analyses revealed that two well-characterized direct VraSR targets, *cwrA* and *vraX* (54, 55), are significantly upregulated under acidic conditions in a VraR-mediated manner, demonstrating the activation of VraSR-mediated response by low pH. In line with this data, highlighting the role of VraSR in acid stress response, recent study has demonstrated that VraSR is essential for the growth of *S. aureus* at low pH (8). Furthermore, in addition to *vraS* and *vraR*, numerous genes encoding proteins involved in cell wall assembly and maintenance were also found to be essential for growth at low pH, thus confirming the importance of cell wall in the acid stress response of *S*. *aureus* (8).

**Figure 7:**
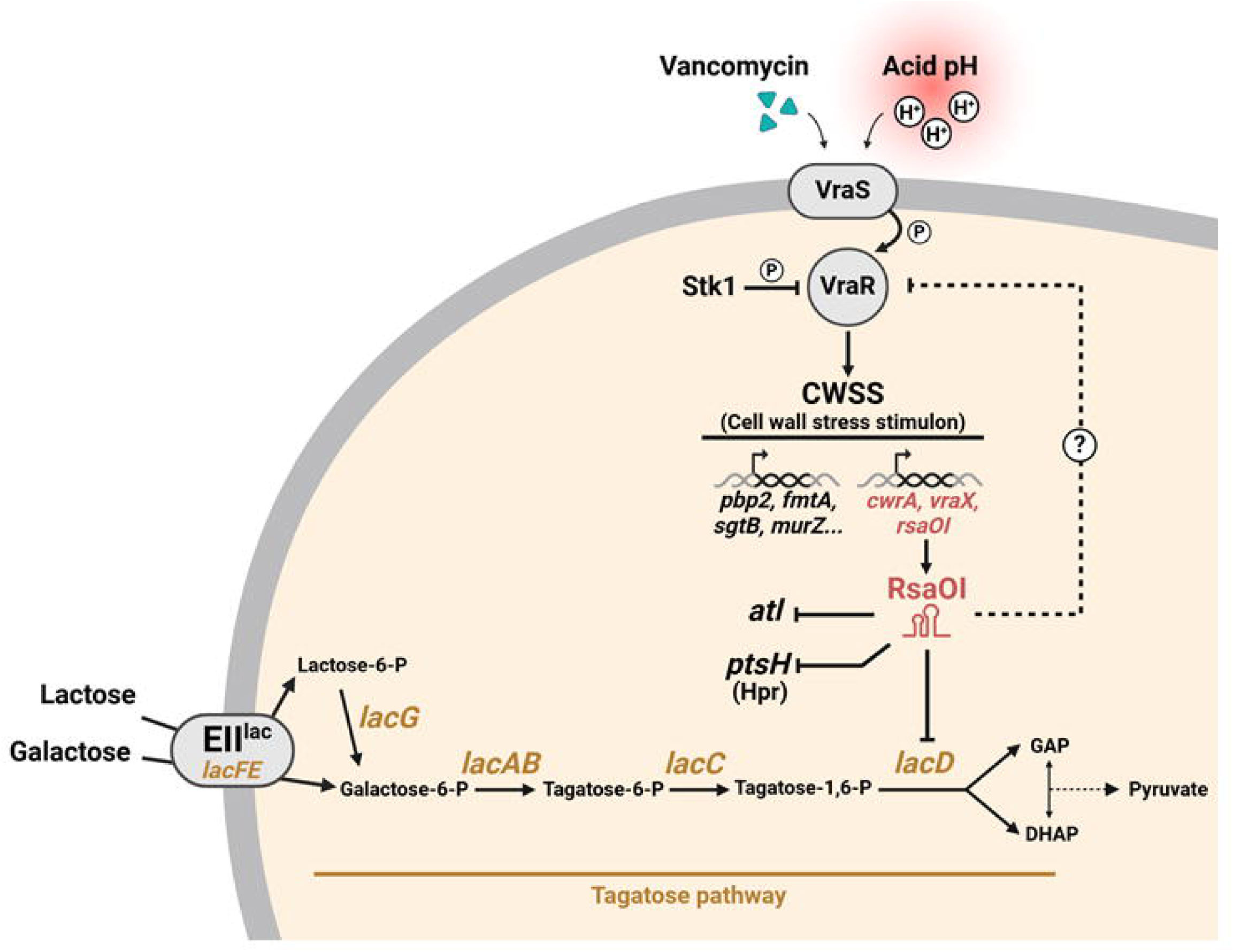
RsaOI acts as a mediator between cell wall stress and the tagatose metabolic pathway. In response to stress generated by exposure to glycopeptides such as vancomycin or to acid stress, the VraSR system, involved in resistance to cell wall-targeting antibiotics, is activated. The sensor unit VraS phosphorylates VraR, thereby activating it, while it was initially kept inactive through phosphorylation by the Stk1 kinase (79). Once activated, VraR induces the transcription of genes belonging to the cell wall stress stimulon (CWSS). This stimulon can be divided into two subsets: the first includes genes induced exclusively upon glycopeptide exposure, primarily involved in cell wall synthesis and dynamics (e.g., *pbp2, fmtA, sgtB, mur*Z); the second subset comprises genes induced both by glycopeptides and acidic conditions (e.g., *cwrA, vraX*, and *rsaOI*, shown in red). Under these stress conditions, RsaOI represses the expression of its target genes: *atl, pts*H (Hpr), and *lacD*. *LacD* encodes the tagatose pathway aldolase, which is essential for galactose metabolism. RsaOI represses *lacD* expression through base pairing at the ribosome-binding site (RBS), thereby reducing tagatose pathway activity and fine-tuning sugar metabolism in response to environmental stressors.

Interestingly, the panel of genes stimulated by VraR varies depending on the perceived stimulus. Notably, acidic stress results in only partial induction of the VraSR-controlled regulon (Figure 7). Our analysis, as well as previously published data on the transcriptional response to acid stress, did not reveal the complete set of VraSR targets that are typically induced in response to cell wall-inhibitory antibiotics (40, 49, 56). The underlying mechanism driving the selective activation of specific genes within the regulon remains unclear, but it could involve differences in VraR phosphorylation dynamics, promoter architecture, or interactions with additional regulatory factors that modulate gene expression in a stimulus-dependent manner. Further studies are needed to elucidate how VraSR coordinates this nuanced regulatory response to diverse environmental challenges.

The identification of *rsaOI* as part of VraR regulon induced by pH lowering aligns with studies showing that expression of *rsaOI* is stimulated by vancomycin treatment (34, 35). Consistent with these findings and the inclusion of *rsaOI* within the cell wall stress regulon, we demonstrated that RsaOI influences bacterial susceptibility to glycopeptide antibiotics. Moreover, the recent study demonstrated that RsaOI controls the expression of the autolysin *atl* involved in *S. aureus* cell division and cell wall turnover, emphasizing the role of RsaOI in cell wall stress response (34). The same study used the CLASH approach to map the sRNA-mRNA interaction network in the Vancomycin-intermediate *S. aureus* (VISA) strain JKD6008 and identified several RsaOI-mRNA interactions associated with the endoribonuclease RNase III (34). In addition to *atl*, RsaOI was found to repress the expression of *ptsH* encoding protein Hpr, involved in catabolite repression by interacting with CcpA (34)(Figure 7). These findings support the role of RsaOI in metabolic regulation and suggest that RsaOI coordinates carbon metabolism and cell wall turnover during vancomycin treatment.

In our study, we analysed the impact of *rsaOI* expression on the protein levels in acidic conditions. Consistent with previous findings (34), we identified that the deletion of *rsaOI* results in an increase in Atl protein levels (Supplementary Table 5), thereby confirming repression of *atl* by RsaOI. However, proteomic analysis of strain in acidic conditions did not reveal changes in Hpr protein levels depending on *rsaOI* expression. It is known that the genetic background of the strains and experimental conditions can significantly influence regulatory network in *S. aureus* and the outcome of sRNA regulation (57, 58). Thus, the differences in RsaOI regulation observed between the two studies could be attributed to distinct stress conditions employed, variations in the genetic backgrounds of used strains as well as their differing levels of glycopeptide susceptibility. In our study, among the proteins whose levels depend on RsaOI, many are involved in carbon metabolism. Proteins such as Fda, SucA and LacD exhibited increased levels in the Δ*rsaOI* strain, suggesting that RsaOI represses their expression. Notably, under our experimental conditions, RsaOI does not affect the levels of Hpr, a protein involved in catabolite repression, indicating that the observed changes in the levels of metabolism-related proteins are independent of Hpr. Altogether, the identification of numerous metabolism-related proteins regulated by RsaOI highlights the sRNA’s pivotal role in metabolic regulation.

It was shown that some sRNA-mRNA interactions recovered by *in vivo* proximity-dependent ligation do not affect mRNA transcript or protein abundance (59). Nevertheless, an mRNAs found to bind RsaOI by RNase III-CLASH analysis was also identified in our study: *rocF* mRNA, which encodes arginase, an enzyme that converts arginine to ornithine (60). Additionally, proteomic analysis revealed that RsaOI reduces HemA protein levels. This finding is consistent with the RNase III-CLASH results, which showed RsaOI binding to *hemX* mRNA, a regulator of *hemA* expression (61). Altogether, RNase III-CLASH capture of RNA-RNA complexes and comparative proteomic analysis under low pH uncovered a broad set of potential RsaOI targets, many of which are linked to cell wall turnover and carbon metabolism. Further studies are needed to elucidate the mechanism of RsaOI action on these targets with connection to cell wall stress.

In this study, the juxtaposition of proteomic data with *in silico* analysis identified *lacD* as a direct target of RsaOI. RsaOI represses the expression of *lacD* by antisense pairing to its ribosome binding site (RBS), resulting in reduced LacD protein production. By downregulating *lacD*, which encodes for tagatose 1,6-diphosphate aldolase, an enzyme of the tagatose pathway, RsaOI modulates galactose metabolism, thereby impacting the bacterium’s capacity to assimilate galactose (Figure 7). Notably, overexpression of *rsaOI* leads to a complete loss of ability to metabolize this sugar.

Bacteria frequently encounter diverse carbon sources in their environments and have developed mechanisms to prioritize the uptake and metabolism of substrates that support rapid growth and competitive advantage. Glucose is the preferred carbon source for most bacteria, which usually suppress the utilization of other sugars in its presence. Furthermore, sugar uptake and metabolism must be carefully regulated to prevent harmful imbalances in metabolite levels and to avoid deleterious accumulation or depletion of metabolites (62, 63). In *S. aureus*, transcription of the *lac* operon is reported to be regulated by multiple mechanisms, including control by the LacR repressor, catabolite repression (64), by glucose in a CcpA-independent manner (65), and it shown to be induced by intracellular galactose-6-phosphate (66). Here we demonstrate that the expression of *lac* operon is also controlled at post-transcriptional level by RsaOI. Similarly, in Gram-negative bacteria, galactose metabolism is controlled not only at the transcriptional level but also post-transcriptionally by sRNA. Spot 42 sRNA binds to the *galETKM* operon mRNA, selectively inhibiting the translation of *galK* and thereby causing discoordinate expression of the *gal* operon (67). This underscores the critical role of post-transcriptional regulation in fine-tuning genes expression for efficient galactose metabolism.

By regulating *lacD*, RsaOI is responsible for the connection between cell wall stress response induced by VraSR and galactose metabolism (Figure 7). In Gram-positive bacteria possessing the Leloir pathway, galactose is metabolized through this pathway leading to the formation of intermediates, UDP-glucose and UDP-galactose, which serve as precursors for cell wall biosynthesis and sugar substitution of lipoteichoic acid (LTA) (68). In *S. aureus*, which does not possess the Leloir pathway, galactose can be converted by the tagatose pathway into tagatose-6-phosphate, which is further processed to produce glyceraldehyde-3-phosphate and dihydroxyacetone phosphate. These intermediates enter central metabolic pathways (like glycolysis), contributing to the pool of building blocks needed for synthesizing essential biomolecules, including the peptidoglycan layers of the cell wall. Disruptions in central metabolism can lead to changes in peptidoglycan synthesis, impacting cell wall thickness and stability (69). Moreover, the enzyme tagatose-1,6-bisphosphate aldolase, encoded by *lacD,* although exhibits its highest affinity for D-tagatose-1,6-bisphosphate but can also use other D-hexose bisphosphate stereoisomers as substrates, including sorbose bisphosphate, psicose bisphosphate, and fructose bisphosphate (70). Fructose monophosphate participates in the formation of UDP-GlcNAc, which is also a precursor for the cell wall (71). Additionally, LacD may possess moonlighting function beyond its role in tagatose pathway, as proteomics studies of *E. coli* identified D-tagatose 1,6-bisphosphate aldolase 2 on the cell surface (72). Interestingly, several studies have reported links between the *lac* operon and cell wall-acting antibiotics, suggesting its potential role in peptidoglycan modification; however, the exact mechanisms remains to be elucidated (73–75).

In conclusion, in *S. aureus*, sRNAs were shown to play a role in the response to variety of environmental stresses, including pH changes, osmotic stress, and nutrient limitations (76). Some of them, such as SprX and 6S RNA, have been found to influence the bacterial susceptibility to antibiotics (27, 77). Here, we report RsaOI as an environment-sensitive sRNA, induced through VraSR two-component system under cell wall stress. RsaOI modulates bacterial metabolism by regulating the tagatose pathway. Therefore, this study highlights the important role of sRNAs in maintaining cellular integrity and the overlapping responses to multiple cell wall-related stress and carbon metabolism.

## ACKNOWLEDGEMENTS

We thank Professor Greg Somerville for staphylococcal strains. We thank Vincent Cattoir for critically reading the manuscript. S.M. thanks the Région Normandie for the engineer financial support. Europe gets involved in Normandy with European Regional Development Fund (ERDF). This project has received funding from the European Union’s Horizon 2020 research and innovation program under the Marie Skłodowska-Curie grant agreement No 101034329. This project has received funding from the Normandy Region under the WINNING Normandy program. MG, HR, AR, and SC thanks Rennes university for financial support (from “Défis émergents” program).

## Author contributions

Supervision: S.C.; Experiments design and Data interpretation : M.G., H.R., K.B.L.H., S.M. N.N., J.H., P.B., A.R., S.C.; Writing – original draft: M.G., and S.C.; Writing – review and editing: M.G., H.R., K.B.L.H., S.M. J.H., P.B., A.R., S.C.; Funding acquisition: A.R., S.C.; Validation: Everyone.

